# Programmable Assembly of DNA Tetrahedra Bearing Atomically Precise Gold Nanoclusters: Stoichiometric Control and Molecular-Level Characterization

**DOI:** 10.64898/2026.06.06.730428

**Authors:** Mathilde Manceau, Abdallah Alhalabi, Christine Saint-Pierre, Elisabetta Boeri-Erba, Xavier Le Guevel, Didier Gasparutto

## Abstract

Atomically precise gold nanoclusters (AuNCs) are ultra-small particles composed of ten to hundreds gold atoms and exhibit unique photophysical properties. Significant progress has been made in tuning and extending their luminescence in the near-infrared window through the design of AuNC assemblies. Herein, we report a straightforward method for synthesizing highly pure, programmable DNA tetrahedra functionalized with a controlled number of AuNCs (from one up to four AuNCs). Using ligand exchange chemistry, AuNCs bearing a single grafted ssDNA onto them were produced. These constructs then served as building blocks for synthesizing tetrahedra through DNA hybridization. Products obtained at each stage of the synthesis were thoroughly characterized using a range of complementary technics. Notably, mass spectrometry in native mode provided novel insights into the accurate composition and stoichiometry of these architectures. This study paves the way for the synthesis and the characterization of a variety of new three-dimensional, DNA-guided AuNC assemblies that may serve as powerful theranostics and biophotonic tools.

## I. Introduction

In the field of nanotechnology, atomically precise gold nanoclusters (AuNCs) are ultra-small particles composed of ten to hundreds gold atoms with core sizes below 2 nm. Their unique properties, such as strong luminescence in the near-infrared window, excellent stability, reduced toxicity and well-defined atomic structures with tunable chemistry makes them promising platforms in biosensing, nanomedicine, and photonics.^1–3^ In recent years, AuNC assemblies have been developed to improve their optical and electronic properties.^4^ Notably, these nanostructures aim to exhibit the aggregation-induced emission (AIE) effect, which enhances their fluorescence properties through spatial confinement.^5,6^ Various materials have been used to guide these assemblies, including polymers, hydrogels and biomolecules.^7–11^

Due to its ability to self-assemble in a highly specific manner, as well as its programmability, biocompatibility and tunability down to 0.3 nm, DNA is of particular interest for guiding AuNCs assemblies.^12,13^ Recently, our group proposed a straightforward method to enable the controlled synthesis of DNA-guided assemblies of atomically precise AuNCs with a mastered interparticle spacing and density.^14^ Two-dimensional AuNC assemblies (dimers, trimers) were obtained with high reproducibility and yields through selective DNA hybridization. However, controlled grafting of atomically precise AuNCs onto three-dimensional DNA structures remains to be achieved. Furthermore, technics for characterizing the composition and stoichiometry of these hybrid architectures are still demanding. Most common approaches provide information about the average size of the nanostructures (*e*.*g*., gel electrophoresis) or their structure (*e*.*g*., microscopy). Determining on the accurate composition and stoichiometry of these architectures would be very helpful. Native mass spectrometry (MS) enables the preservation of non-covalent interactions to study macromolecular assemblies. In particular, native MS assesses the mass of intact non-covalent complexes to determine their stoichiometry and identify direct interactions between their components.

The aim of this study is to address these gaps by synthesizing programmable DNA tetrahedra functionalized with a controlled number (one, two, three or four) of atomically precise AuNCs. The know structure Au_25_SR_18_ (SR=para-mercaptobenzoic acid, pMBA) was used as archetype.^15–18^ Numerous analytical methods were employed to thoroughly analyze the products obtained at each step of the synthesis process. In addition, the final biohybrid nanoarchitectures were characterized using complementary native methods. Gel electrophoresis, dynamic light scattering (DLS), size exclusion chromatography (SEC), native MS and microscopy confirmed their integrity and purity.

## II. Materials and methods

### II.1. Chemicals

Acrylamide solution (40 % 19:1 acrylamide/bis(acrylamide)), triethylammonium acetate (TEAA) 1 M, ammonium acetate 7.5 M, TBE buffer 10X molecular biology grade and agarose were purchased from Sigma-Aldrich (USA). Acetonitrile (ACN) and tris(2-carboxyethyl)phosphine hydrochloride (TCEP) were from Fisher Scientific (UK) and Thermo Scientific (USA), respectively. Glycerol was purchased from AppliChem (Germany) and absolute denatured ethanol from Carlo Erba Reagents (France). SYBR Gold (10000X in dimethyl sulfoxide) was from Thermo Fisher Scientific (USA). Oligonucleotides (ODNs) were ordered from Biomers (Germany). Their sequences and the structure of the 5’-thiol phosphoramidite monomer are reported in Table S1. Chemical products for AuNC synthesis were purchased from Sigma-Aldrich (France).

### II.2. Au_25_pMBA_18_ synthesis and purification

Au_25_pMBA_18_ clusters were prepared and purified according to the procedure described in our previous study.^14^ Briefly, HAuCl_4_ (2.54 mmol, 100 mg) was added to a 116 mM para-mercaptobenzoic acid (pMBA) solution composed of 40 mL of methanol and 4 mL of tributylamine. After stirring for 30 minutes at room temperature, a gold thiolate/tributylamine complex was formed. Then, 200 mg of trimethylamine borane was added with stirring for 2 hours to induce slow gold reduction, followed by the addition of a further 200 mg. The clusters were obtained by stirring for 24 hours overnight. After several dissolution/precipitation cycles, the crude solution was purified using native PAGE with a 3 % stacking gel and a 20 % separating gel containing a 19:1 acrylamide/bis(acrylamide) ratio. The corresponding Au_25_pMBA_18_ band was collected and crushed in a 2 mL Ultrafree centrifugal filter (Merck). 200 µL of a 0.1 M triethylamine acetate (TEAA) + 5 % acetonitrile (ACN) solution was added to the filter. The filter was then vortexed and centrifuged at 12000 g for 10 minutes. This step was repeated until the majority of the Au_25_pMBA_18_ clusters had been recovered. Au_25_pMBA_18_ concentration was determined by UV spectroscopy at 700 nm with an extinction coefficient of 6997 M^-1^.cm^-1^. This solution was stored at 4°C until use.

### II.3. Synthesis of Au_25_pMBA_(18-n)_SSX_(n)_ bricks

Au_25_pMBA_18_ clusters were functionalized with 55-mer oligonucleotides (SS1, SS2, SS3 or SS4) that were modified with a HS-(CH_2_)_6_ linker at their 5’ end. Prior to grafting, the ODNs were reduced with tris(2-carboxyethyl)phosphine hydrochloride (TCEP) and purified by ethanol precipitation. The reduced ODNs were then mixed with three times more nanomoles of the Au_25_pMBA_18_ clusters (typically 6 and 18 nmoles, respectively), and 7.5 M ammonium acetate was added to reach a final concentration of 300 mM. The solution was then incubated in an Eppendorf Thermomixer at 40 °C with stirring at 450 rpm overnight. The next morning, the solution was allowed to cool to room temperature before being stored at -20 °C.

### II.4. Purification of Au_25_pMBA_17_SSX_1_ bricks by PolyAcrylamide Gel Electrophoresis (PAGE)

Native PAGE was carried out using a vertical gel electrophoresis unit. The separating gel was prepared with a final concentration of 20 % 19:1 acrylamide/bis(acrylamide) and the stacking gel with 3 % concentration. In the meantime, the crude solution of Au_25_pMBA_(18-n)_SSX_(n)_ bricks was defrost and concentrated using a 3 kDa centrifugal concentrator (Amicon 500, Merck) at 14000 g for 20 minutes after addition of 5 % ACN to avoid nonspecific adsorption of the nanoclusters on the membrane. The recovered volume was adjusted to 50 µL and 25 µL of 50 % glycerol aqueous solution was added to the solution. The resulting 75 µL were loaded on the gel and migration was done using TBE 1X buffer (89 mM Tris, 89 mM boric acid, 2 mM EDTA) at 140 V for around 2 hours. The corresponding Au_25_pMBA_17_SSX_1_ band was collected and crushed in a 2 mL Ultrafree centrifugal filter (Merck). 200 µL of a 0.1 M triethylammonium acetate + 5 % ACN solution was added to the filter. The filter was then vortexed and centrifuged at 12000 g for 6 minutes. This step was repeated until the gel mixture was colorless, meaning that the majority of the Au_25_pMBA_17_SSX_1_ had been recovered.

### II.5. Self-assembly of the functionalized DNA tetrahedra

600 pmol of each of the four complementary ODN sequences SSX, with or without a gold nanocluster, were mixed in a 6 mL of 0.3 M TEAA solution. The temperature was increased to 75 °C for 15 minutes, after which the solution was allowed to cool to room temperature slowly overnight. The resulting tetrahedron was then filtered and concentrated three times using a 50 kDa centrifugal concentrator (Amicon 4, Merck) at 7500 g for 5 minutes. The concentration of the tetrahedron was estimated by UV spectroscopy using the tetrahedron extinction coefficient at 260 nm of 1.87×10^6^ M^-1^.cm^-1^,^19^ assuming that the presence of gold nanocluster did not significantly impact on its absorbance at this wavelength (Au_25_pMBA_18_ extinction coefficient at 260 nm of 1.5×10^5^ M^-1^.cm^-1^).

### II.6. Analytical PolyAcrylamide Gel Electrophoresis (PAGE)

Native or denaturing PAGE were carried out using a vertical gel electrophoresis unit with a two gels holder. Native gels were prepared with a final concentration of 20 % 19:1 acrylamide/bis(acrylamide). Denaturing gels were prepared with a final concentration of 15 % 19:1 acrylamide/bis(acrylamide) with 7 M urea. For DNA visualization, each of the solutions deposited in the wells of the gel was composed of 100 ng of oligonucleotide in TBE 1X, 1 µL of the intercalating agent (SYBR Gold diluted by 10^3^ in TBE 1X), 2 µL of the loading buffer (DNA loading buffer 6X, Lonza) and a sufficient amount of TBE 1X to reach a final volume of 10 µL. For nanoclusters visualization, each of the solutions deposited in the wells of the second gel was composed of 3 µg of oligonucleotide in TBE 1X, 2 µL of the loading buffer (50 % glycerol in TBE 1X) and a sufficient amount of TBE 1X to reach a final volume of 10 µL. The migration was conducted at 140 V for approximately 2 hours for native gel and 350 V for denaturing gel. The gels were imaged either by an ultraviolet gel documentation system (Gel Doc XR+, BioRad, USA) for DNA visualization or in the shortwave infrared optical window (SWIR; 900 – 1700 nm) using a NIR-II Princeton camera 640ST coupled with a laser excitation source at 808 nm (100 mW.cm^−2^) for nanoclusters visualization. A short-pass excitation filter (Thorlabs) was used at 1000 nm and a long-pass emission filter (LP) set at 1064 nm (Semrock) was coupled onto a 25 mm lens with a numerical aperture of 4 (Navitar).

### II.7. Agarose Gel Electrophoresis (AGE)

AGE was conducted using a horizontal gel electrophoresis unit. The separating gel was composed of 150 mL of 3 % agarose in 0.5X TBE. Each of the solutions deposited in the wells was composed of 100 ng of oligonucleotide in TBE 0.5X, 1 µL of the intercalating agent (SYBR Gold diluted by 10^3^ in TBE 0.5X), 2 µL of the loading buffer (DNA loading buffer 6X, Lonza) and a sufficient amount of TBE 0.5X to reach a final volume of 10 µL. The migration was conducted at 130 V for approximately 2 hours. The gel was imaged by an ultraviolet gel documentation system (Gel Doc XR+, BioRad, USA).

### II.8. Size Exclusion Chromatography (SEC)

Size Exclusion Chromatography (1100 Series HPLC system, Agilent technologies) was used to analyze the synthetized Au_25_pMBA_17_SSX_1_ bricks after their purification by native PAGE. Samples were eluted using a XBridge Premier Protein SEC column (250 Å, 2.5 µm, 4.6×300 mm, Waters) at 30 °C. The mobile phase was 0.1 M TEAA/ACN 95/5 (v/v) and the flow rate was set to 200 µL/min. 50 µL containing 50 pmol of the samples were injected on the column and their detection was performed by UV-Visible absorbance measurement at 260 nm and 350 nm using a diode array detector.

### II.9. Native Mass Spectrometry (MS)

Before native MS, buffer of Au_25_pMBA_17_SSX_1_ bricks and DNA tetrahedra was exchange by 0.3 M ammonium acetate using 10 kDa centrifugal concentrator (Vivaspin 500, Merck) at 12000 g for 30 minutes after addition of 5 % ACN to avoid nonspecific adsorption or 50 kDa centrifugal concentrator (Amicon 4, Merck) at 7500 g for 5 minutes, respectively. After their UV adsorption at 260 nm measurement, a sufficient amount of 0.3 M ammonium acetate was added to the solutions to reach a final concentration ranging between 5 and 10 µM.

The samples were analyzed by native mass spectrometry.^20^ Protein ions were generated using a nanoflow electrospray (nano-ESI) source. Nanoflow platinum-coated borosilicate electrospray capillaries were bought from Thermo Electron SAS (Courtaboeuf, France). MS analyses were carried out on a quadrupole time-of-flight mass spectrometer (Q-TOF Ultima, Waters Corporation, Manchester, U.K.). The instrument was modified for the detection of high masses.^21,22^ The following instrumental parameters were used: capillary voltage = 1.2 – 1.3 kV, cone potential = 40 V, RF lens-1 potential = 40 V, RF lens-2 potential = 1 V, aperture-1 potential = 0 V, collision energy = 30 – 100 V, and microchannel plate (MCP) = 2100 V. All mass spectra were calibrated externally using a solution of cesium iodide (6 mg/mL in 50 % isopropanol) and were processed using the Masslynx 4.0 software (Waters Corporation, Manchester, U.K.), Massign software package^23^ and UniDec.^24^

### II.10. Dynamic Light Scattering (DLS)

DLS (Litesizer 500, Anton Paar) was used to obtain the number distribution curves for the tetrahedral solutions. Measurements were performed in triplicate at a side scatter angle of 90° using a low-volume quartz cell, with tetrahedron concentrations ranging from 12.0 to 23.7 µM and volumes ranging from 25 to 35 µL. Data were considered valid if the transmittance was greater than 70 % and g1^2^ was between 0.5 and 0.9.

### II.11. Transmission electron microscopy (TEM)

TEM formvar grids were first pre-coated with a polylysine solution (0.1 % in water) for 5 min and then washed twice with deionized water to provide a positively charged surface layer, enabling electrostatic interaction with the samples. Diluted AuNC-DNA tetrahedra (1-2 µM) were deposited onto the pre-coated TEM grids for 30 min and excess solution was removed by capillary action using tissue. Morphologies of the AuNC-DNA tetrahedra samples were examined by TEM using a FEG JEOL 2100F system operating at 200 kV. Energy-dispersive X-ray (EDX) measurements were performed to confirm the presence of gold within the self-assemblies.

## III. Results and discussion

### III.1. Au_25_pMBA_17_SSX_1_ constructs as self-assembly building blocks

Due to its widespread use in biomedical applications such as biosensing and drug delivery, DNA tetrahedron was selected as the three-dimensional model for investigating stoichiometrically controlled gold nanocluster (AuNC) grafting.^25,26^ The tetrahedral construct was composed of four single-stranded DNA (ssDNA), namely SS1, SS2, SS3 and SS4, each of which containing 55 nucleotides. These strands were chosen for their ability to hybridize and form a tetrahedron with 17-base-long sides (Figure 1A).^27^ For the sake of clarity, the strands are henceforth referred as SSX. Our strategy requires two main steps: (i) grafting of a single 5’-thiolated DNA strand on a single AuNC via ligand exchange, forming Au_25_pMBA_17_SSX_1_ constructs (Figure 1B), and (ii) hybridization of relevant strands forming tetrahedra with zero, one, two, three or four AuNCs, named TDN 0, TDN 1, TDN 2, TDN 3 and TDN 4 respectively (Figure 1C).

**Figure 1.**
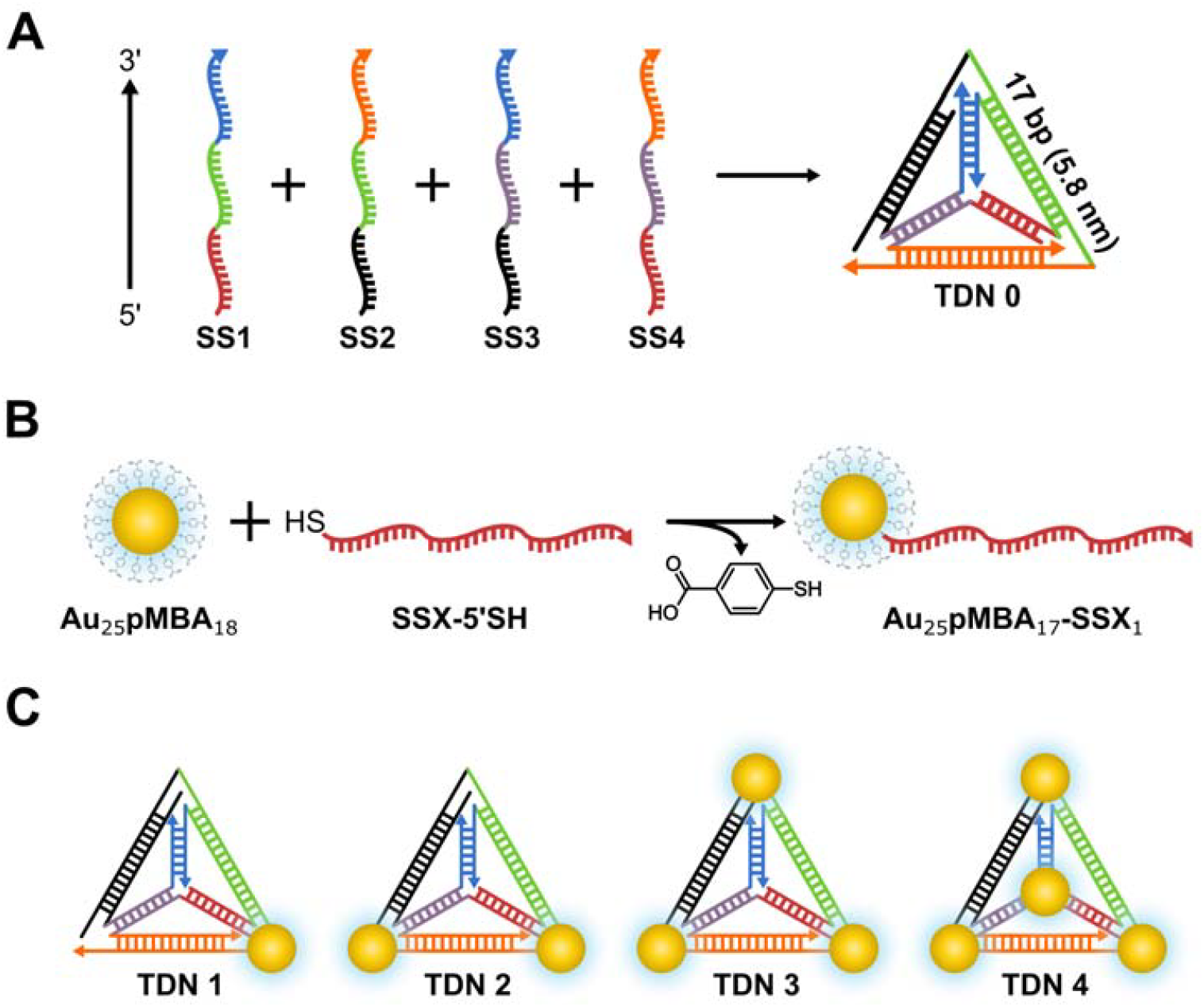
Functionalization strategy of DNA tetrahedra with gold nanoclusters (AuNCs). Representation of the self-assembly of the four complementary sequences (SS1, SS2, SS3 and SS4) into a tetrahedron (TDN) (A), the ligand exchange reaction of a thiolated ssDNA with a pMBA ligand of the gold nanocluster (B), and the different tetrahedra functionalized with a controlled number of AuNCs to be obtained (C).

A three-fold excess of Au_25_pMBA_18_ was used for direct ligand exchange of thiolated SSX to promote the grafting of a single oligonucleotide (ODN) onto AuNCs. Indeed, we have recently demonstrated that such a ratio could enhance monosubstituted AuNC-ODN constructs yields.^14^ The Au_25_pMBA_17_SSX_1_ constructs were subsequently purified from Au_25_pMBA_18_ in excess and Au_25_pMBA_16_SSX_2_ by PAGE with yield ranging from 26 % to 37 % (Figure S1). To evaluate the efficiency of this purification process, analytical denaturing (Figure 2A) and native (Figure S2) PAGE were conducted. Both gels revealed a unique band for the purified Au_25_pMBA_17_SSX_1_ using SYBR Gold staining, with a slower migration than SSX oligonucleotides. Visualization of the native gel under brightfield showed that the ungrafted AuNCs eluted faster than the DNA strands due to their smaller size (Figure S2B). Furthermore, the same samples were deposited on another native polyacrylamide gel at a higher concentration. Visualization of the gel under the SWIR window showed that the purified bands contained AuNCs (Figure 2B). These results are consistent with previous literature and suggest that pure Au_25_pMBA_17_SSX_1_ constructs were obtained.^14^

**Figure 2.**
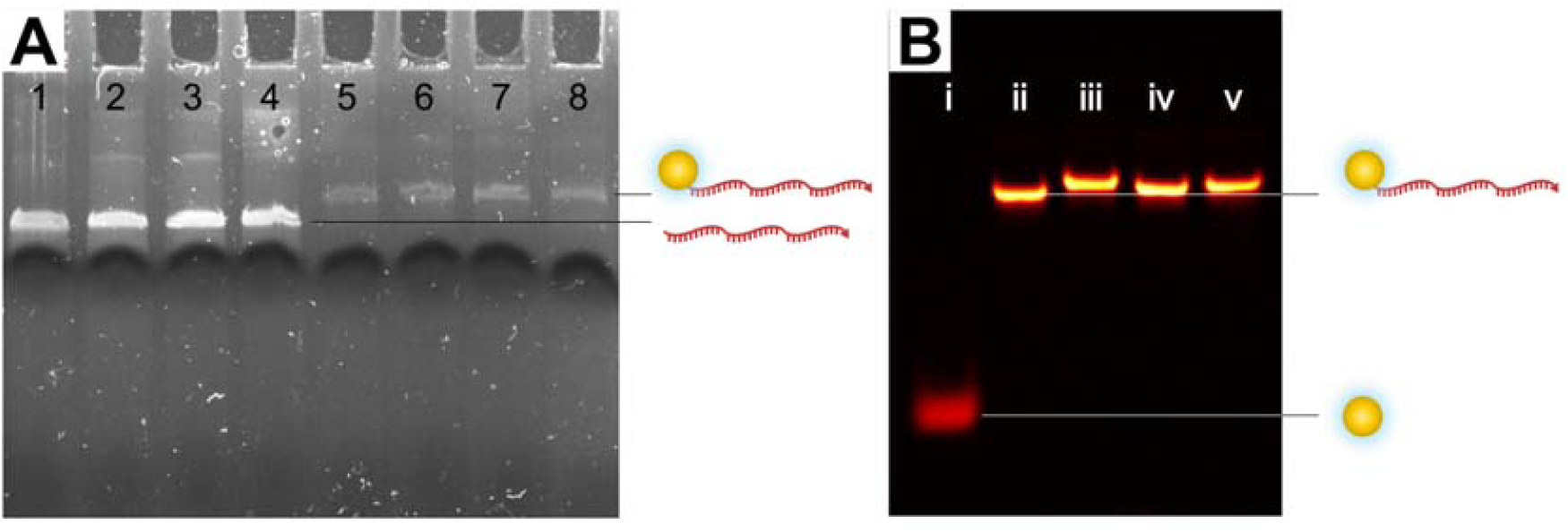
15 % denaturing 19:1 polyacrylamide gel electrophoresis (PAGE) analysis of the purified Au_25_pMBA_17_SSX_1_ under UV light after SYBR gold staining (A): lane 1, SS1; lane 2, SS2; lane 3, SS3; lane 4, SS4, lane 5, Au_25_pMBA_17_SS1_1_; lane 6, Au_25_pMBA_17_SS2_1_; lane 7, Au_25_pMBA_17_SS3_1_; lane 8, Au_25_pMBA_17_SS4_1_. Bands of Au_25_pMBA_17_SSX_1_ are less intense due to a quenching effect of the SYBR Gold stain. 20 % native 19:1 PAGE analysis of the purified Au_25_pMBA_17_SSX_1_ under fluorescence with an excitation at 808 nm and emission at 950 nm (B): lane i, Au_25_pMBA_18_; lane ii, Au_25_pMBA_17_SS1_1_; lane iii, Au_25_pMBA_17_SS2_1_; lane iv, Au_25_pMBA_17_SS3_1_; lane v, Au_25_pMBA_17_SS4_1_.

These products were further analyzed by UV absorbance spectroscopy which showed that Au_25_pMBA_17_SSX_1_ spectra contained the characteristic bands of both Au_25_pMBA_18_ (350, 430, 480 and 690 nm) and DNA (260 nm) (Figure 3A). The samples exhibited similar photoluminescence (PL) profiles when excited at 690 nm and 808 nm (Figure S3). The area under the PL emission peaks was corrected in relation to the absorbance of the samples at 690 nm. Au_25_pMBA_17_SSX_1_ have PL emission 1.8 ± 0.4 larger than Au_25_pMBA_18_ (Table S2). This could be explained by the presence of ssDNA which is a rich electron donor^28^ but also to the flexibility of the ligand protecting the AuNC core from the environment.^17,29^ The elution times of Au_25_pMBA_18_, SSX and Au_25_pMBA_17_SSX_1_ were compared using size exclusion chromatography (SEC) coupled with absorbance spectroscopy (Figure 3B, Figure S4). ODNs absorb four times more at 260 nm than AuNCs (extinction coefficients around 6.0×10^5^ M^-1^.cm^-1^ and 1.5×10^5^ M^-1^.cm^-1^, respectively) and do not absorb at 350 nm. Therefore, the presence of ODN and AuNCs in the analyzed samples could be tracked by monitoring the absorption at these two wavelengths. The resulting chromatograms show similar elution times for all the SSX strands and the Au_25_pMBA_17_SSX_1_ constructs. Au_25_pMBA_18_ nanoclusters (16.6 min) have a longer elution time than SSX (12.5 min) and Au_25_pMBA_17_SSX_1_ (12.1 min) have a slightly longer elution time than SSX. This is consistent with expectations, since in SEC, samples elute according to their hydrodynamic volume, with larger particles eluting faster than smaller ones. Purity of the Au_25_pMBA_17_SSX_1_ constructs can be checked using this method. The presence of a single peak with strong absorption at 260 nm and a weaker absorption at 350 nm indicates that the samples are pure and contain both AuNCs and ODN. It can be noted that the peaks show a tailing asymmetry, a common feature of SEC that reveals interactions between the samples and the stationary phase, such as hydrophobic interactions.^30^

**Figure 3.**
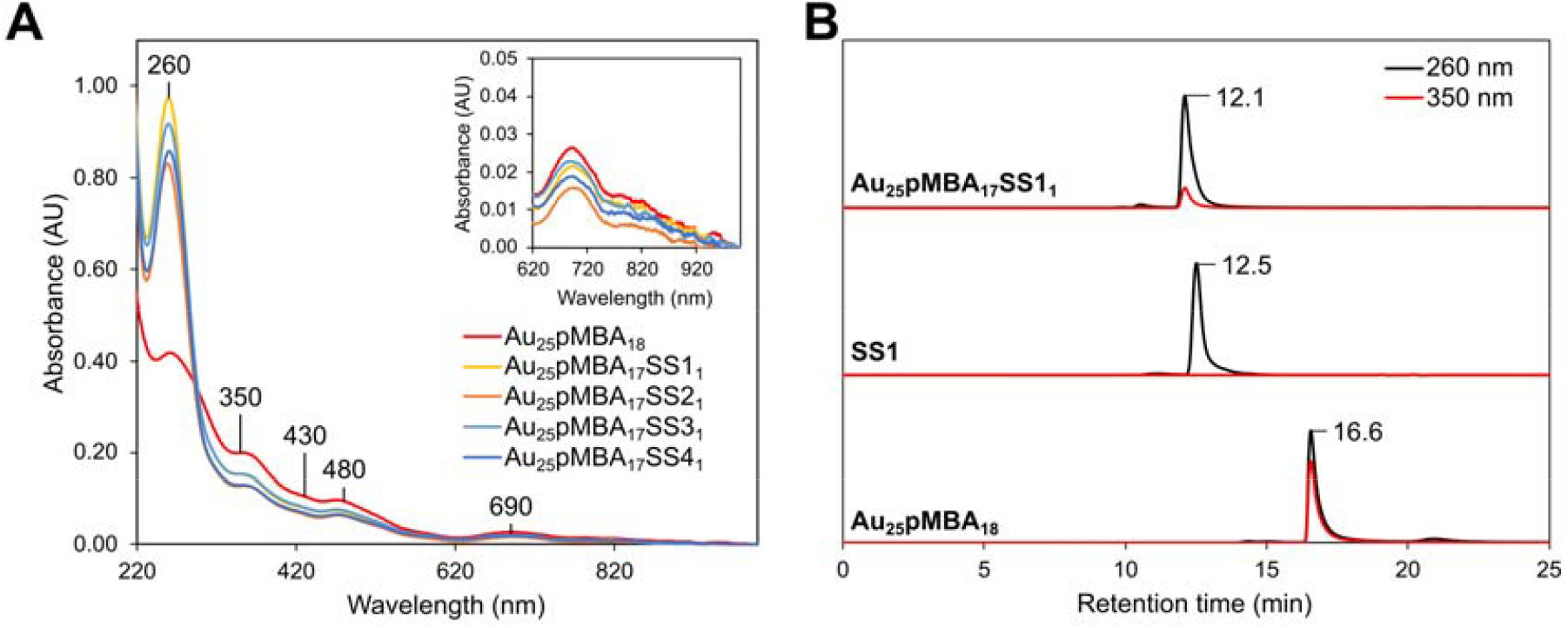
UV-Vis absorbance spectra of the four different Au_25_pMBA_17_SSX_1_ and Au_25_pMBA_18_ nanoclusters (A). Au_25_pMBA_17_SSX_1_ spectra contain the characteristic bands of both Au_25_pMBA_18_ (350, 430, 480 and 690 nm) and DNA (260 nm). Size exclusion chromatograms of Au_25_pMBA_17_SS1_1_, SS1 and Au_25_pMBA_18_ (B) with UV detection at 260 nm (black) and 350 nm (red).

To gain accurate insight into the composition of these constructs, native MS was used (Figure 4). The measurements in positive mode showed three main charge states of Au_25_pMBA_17_SSX_1_ (*i*.*e*., from 6+ to 8+). The *m/z* values and charge states (*z*) of each compound are listed in Table S3. The resulting experimental masses (24603 ± 1 Da; 24754 ± 2 Da; 24635 ± 2 Da and 24615 ± 3 Da) were in agreement with the theoretical masses (24603 Da; 24755 Da; 24633 Da; 24612 Da, respectively). It can be concluded that Au_25_pMBA_17_SSX_1_ constructs were successfully obtained with a high degree of purity. The MS investigation of Au_25_pMBA_17_SS2_1_ revealed the presence of gas phase dimers (Figure 4B). This phenomenon may be attributed to the stacking of the SS2 strand. Consistently, the SS2 strand alone was observed to form dimers (Figure S5).

**Figure 4.**
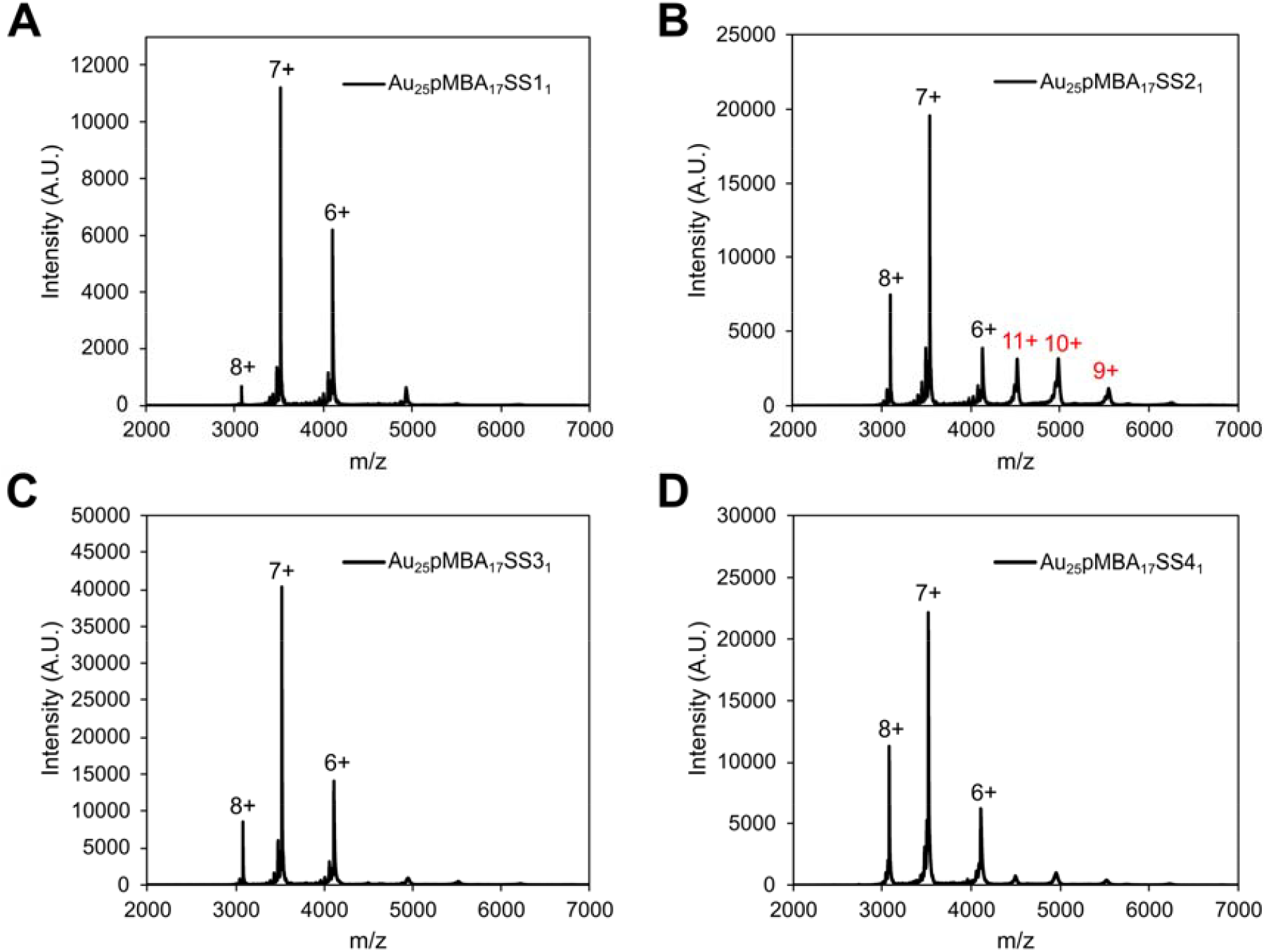
Native MS spectra of Au_25_pMBA_17_SS1_1_ (A), Au_25_pMBA_17_SS2_1_ (B), Au_25_pMBA_17_SS3_1_ (C), and Au_25_pMBA_17_SS4_1_ (D) bricks. The different charge states of Au_25_pMBA_17_SSX_1_ ions are represented in black while the charge states of Au_25_pMBA_17_SS2_1_ dimer are in red.

### III.2. Self-assembly of DNA tetrahedra with programmable Au_25_pMBA_18_ incorporation

Following the successful purification of Au_25_pMBA_17_SSX_1_ constructs, tetrahedra containing zero, one, two, three or four AuNCs (TDN 0, TDN 1, TDN 2, TDN 3 and TDN 4, respectively) could be synthetized. For this, the four oligonucleotide strands (SS1, SS2, SS3 and SS4), with or without an AuNC depending on the desired tetrahedron, were hybridized. The resulting products, with yields ranging from 67 % to 77 %, were analyzed using UV-visible spectroscopy. (Figure 5A). After normalization at 260 nm, the TDN 0 spectrum contained only the characteristic DNA band (absorption at 260 nm). By contrast, the TDN 1, TDN 2, TDN 3 and TDN 4 spectra contained the characteristic bands of both Au_25_pMBA_18_ (350, 430, 480 and 690 nm) and DNA (260 nm). Furthermore, the intensity of the Au_25_pMBA_18_ AuNCs characteristic band (690 nm) increased linearly from TDN 1 to TDN 4 (Figure S6).^15,31^ Consequently, TDN 4 appeared to contain four more AuNCs than TDN 1. No shift of the 690 nm absorption band was observed on any of the AuNC-labeled-TDN. Moreover, the AuNCs increase from TDN 0 to TDN 4 was also visible to the naked eye, as the intensity of the brown-orange color increased (Figure S7). When excited at 690 nm and 808 nm, the photoluminescence (PL) emission band record of the tetrahedra containing AuNCs showed the same PL profile. This consists of at least three bands centered at 930 nm, 1090 nm and 1250 nm (Figure 5B, Figure S8), as already reported for Au_25_SR_18_ species such as Au_25_pMBA_18_.^1,14,32,33^ As previously, the area under the curve of the PL emission peaks were corrected in relation to the concentration of the tetrahedra. After correction, the intensity of the PL emission was found to be proportional to the number of AuNCs on the tetrahedra (Figure 5B inset). The absence of significant changes in PL properties can be explained by the distance between each AuNC being superior to 5 nm. Thus, AuNCs are too distant to allow aggregation-induced enhancement (AIE) of fluorescence.^34^ Additionally, these tetrahedra are relatively flexible, and the limited rigidity of such nanostructures may also account for the absence of PL enhancement. Both UV-visible and fluorescence spectroscopies yielded results that are in agreement with the correct synthesis of the tetrahedra. However, it should be noted that these techniques provide an average result influenced by the content of the entire solution. Therefore, they cannot provide accurate information on the purity of the synthesized tetrahedra. Dynamic light scattering (DLS) measurements were performed in triplicate. The number distribution curves showed a single narrow peak indicating homogeneous solutions, with maximum values of 9.18 ± 0.42 nm, 8.24 ± 2.25 nm, 8.26 ± 0.75 nm, 7.44 ± 1.00 nm and 7.81 ± 0.36 nm from TDNs 0 to 4, respectively (Figure S9). Consequently, TDNs could not be sorted by size using DLS. This could be explained by the AuNCs being too small for DLS resolution. TEM imaging further confirmed the presence of nanostructures ranging from 10 to 30 nm, with gold identified by EDX analysis (Figure S10). However, imaging the internal organization of the AuNCs within these assemblies accurately proved difficult. Indeed, the tetrahedra degraded under the electron beam at high magnification despite multiple attempts. Further optimization of sample preparation (solution, staining, cross-linking, cryo-TEM…) and imaging conditions would be required to achieve high-resolution TEM characterization of these assemblies at the nanometer scale.

**Figure 5.**
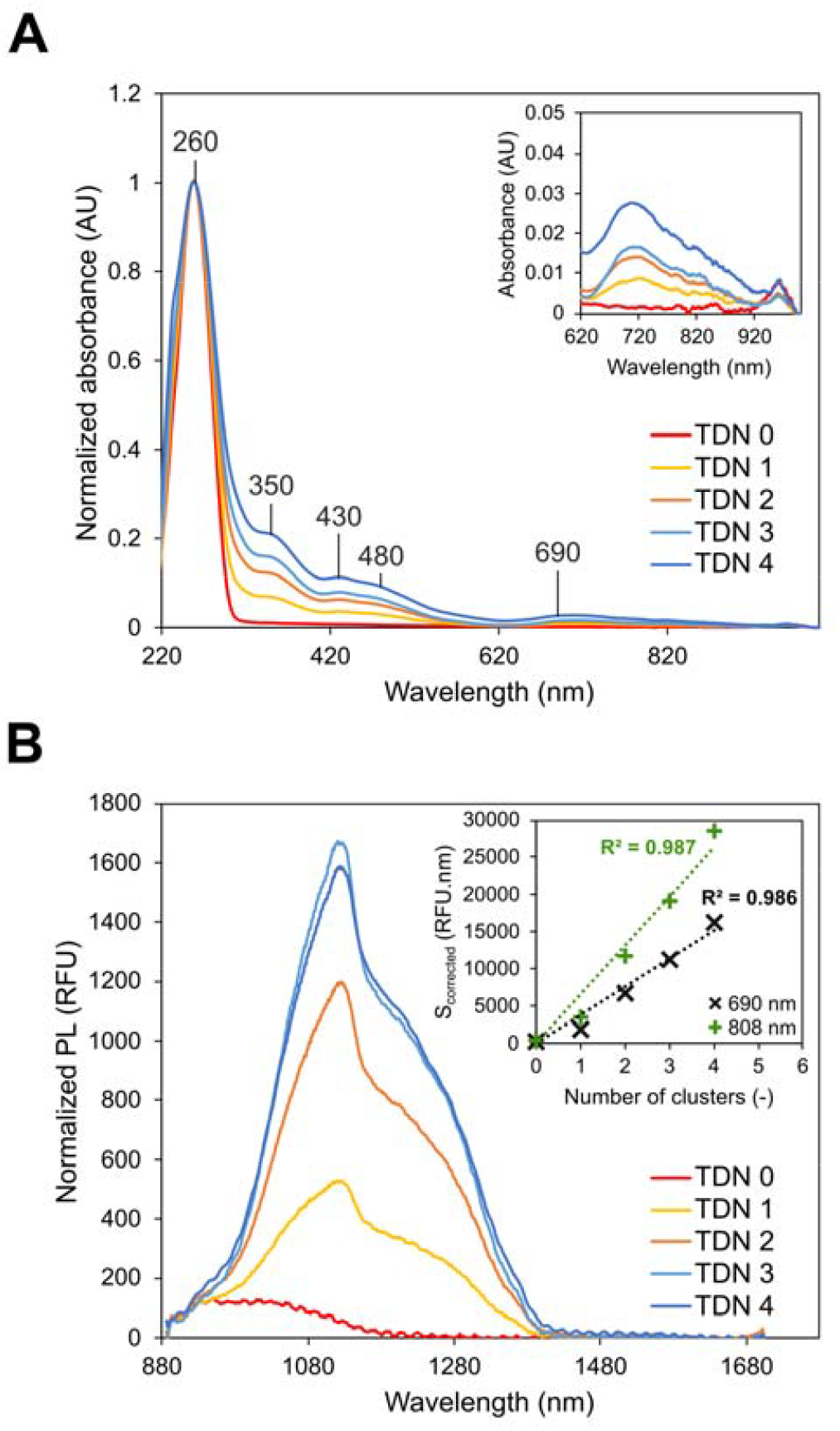
UV-Vis absorbance spectra of the different tetrahedra (A). TDN 1, TDN 2, TDN 3 and TDN 4 spectra contain the characteristic bands of both Au_25_pMBA_18_ (350, 430, 480 and 690 nm) and DNA (260 nm). By contrast, TDN 0 spectrum shows only DNA band. Photoluminescence (PL) emission peaks of the tetrahedra with excitation at 690 nm (B). The inset represents the corrected area of the PL emission peaks against the number of gold nanoclusters contained in the tetrahedra, with excitation at 690 nm (black) or 808 nm (green).

The purity of the tetrahedra was investigated using agarose gel electrophoresis (AGE). The gel revealed unique bands for each tetrahedron (Figure 6). Their order of migration, from fastest to slowest, was as follows: TDN 0 > TDN 1 > TDN 2 > TDN 3 > TDN 4. This is consistent with the slower migration of ssDNA after grafting onto an AuNC (lanes 2 and 3, Figure 6). Overall, the tetrahedra appeared to be pure, and their migration was aligned with their theoretical number of AuNCs.

**Figure 6.**
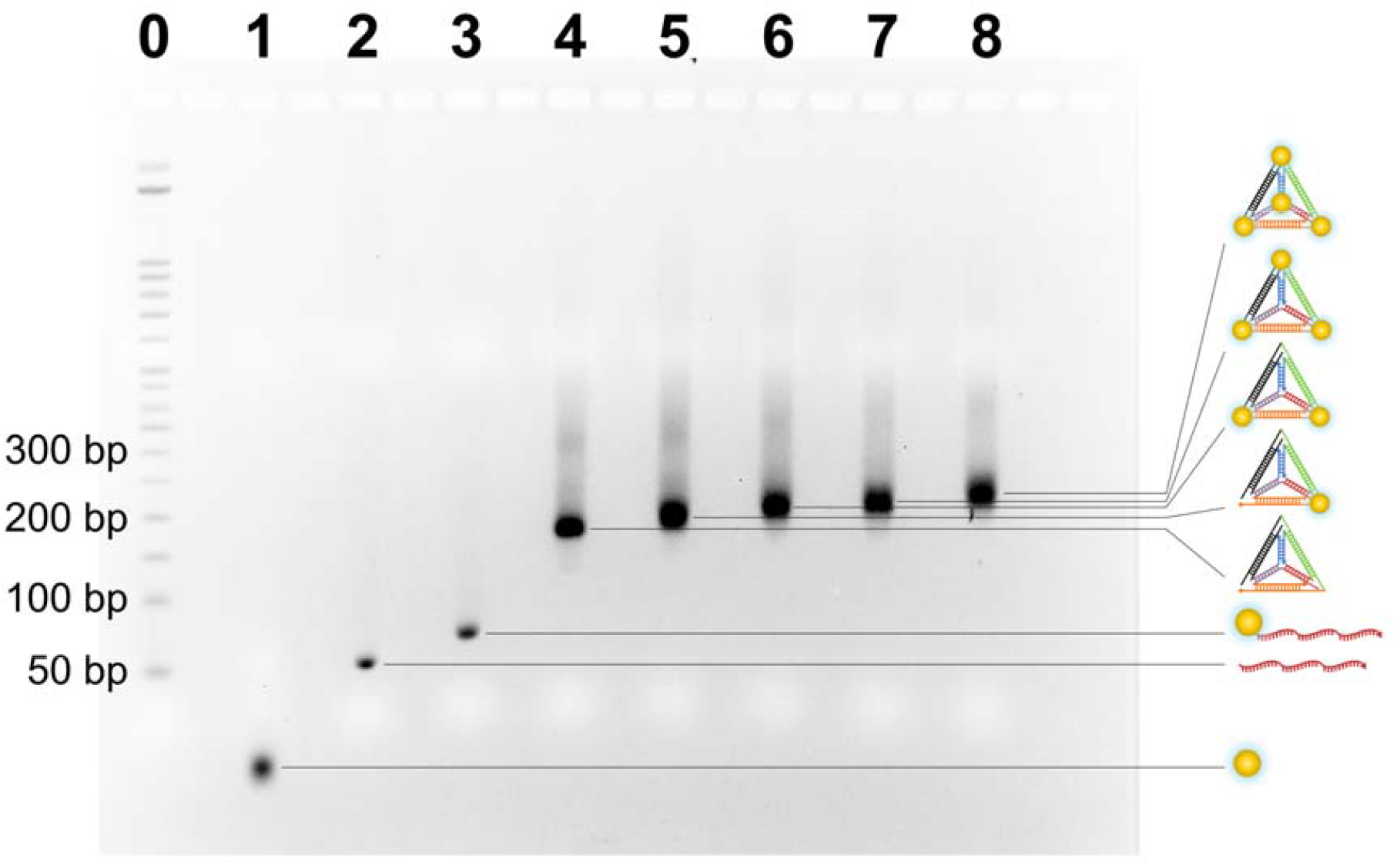
Agarose gel electrophoresis (3% AGE) analysis of the resulting tetrahedra under UV light after SYBR gold staining: lane 0, 50 bp DNA step ladder; lane 1, Au_25_pMBA_18_; lane 2, SS1; lane 3, Au_25_pMBA_17_SS1_1_; lane 4, TDN 0; lane 5, TDN 1; lane 6, TDN 3; lane 7, TDN 3; lane 8, TDN 4.

The tetrahedra were further analyzed by native MS (Figure 7). This work provides a new demonstration of the characterization of DNA tetrahedra with atomically precise metal nanoclusters by mass spectrometry. Measurements in positive mode showed different charge states ranging from 11+ to 17+ depending on the tetrahedron. The *m/z* values of each compound’s charge states are in Table S4. Each charge state 14+ was shifted by a Δ*m/z* of approximately 552, corresponding to a mass of ―, *i*.*e*., the addition of a AuNC with the thiol function of a ssDNA (7723 Da). The resulting experimental masses of TDN 0, TDN 1, TDN 2, TDN 3 and TDN 4 were determined (67712 ± 2 Da; 75436 ± 7 Da; 83163 ± 5 Da; 90881 ± 7 Da and 98599 ± 6 Da, respectively). These are consistent with the theoretical masses of the tetrahedra (67711 Da; 75434 Da; 83158 Da; 90881 Da; 98604 Da, respectively). The tetrahedra were determined to be of a high degree of purity, thus validating the efficiency of the assembly and purification processes. Overall, native MS represents a powerful approach for analyzing the integrity and purity of a DNA-AuNC architectures.

**Figure 7.**
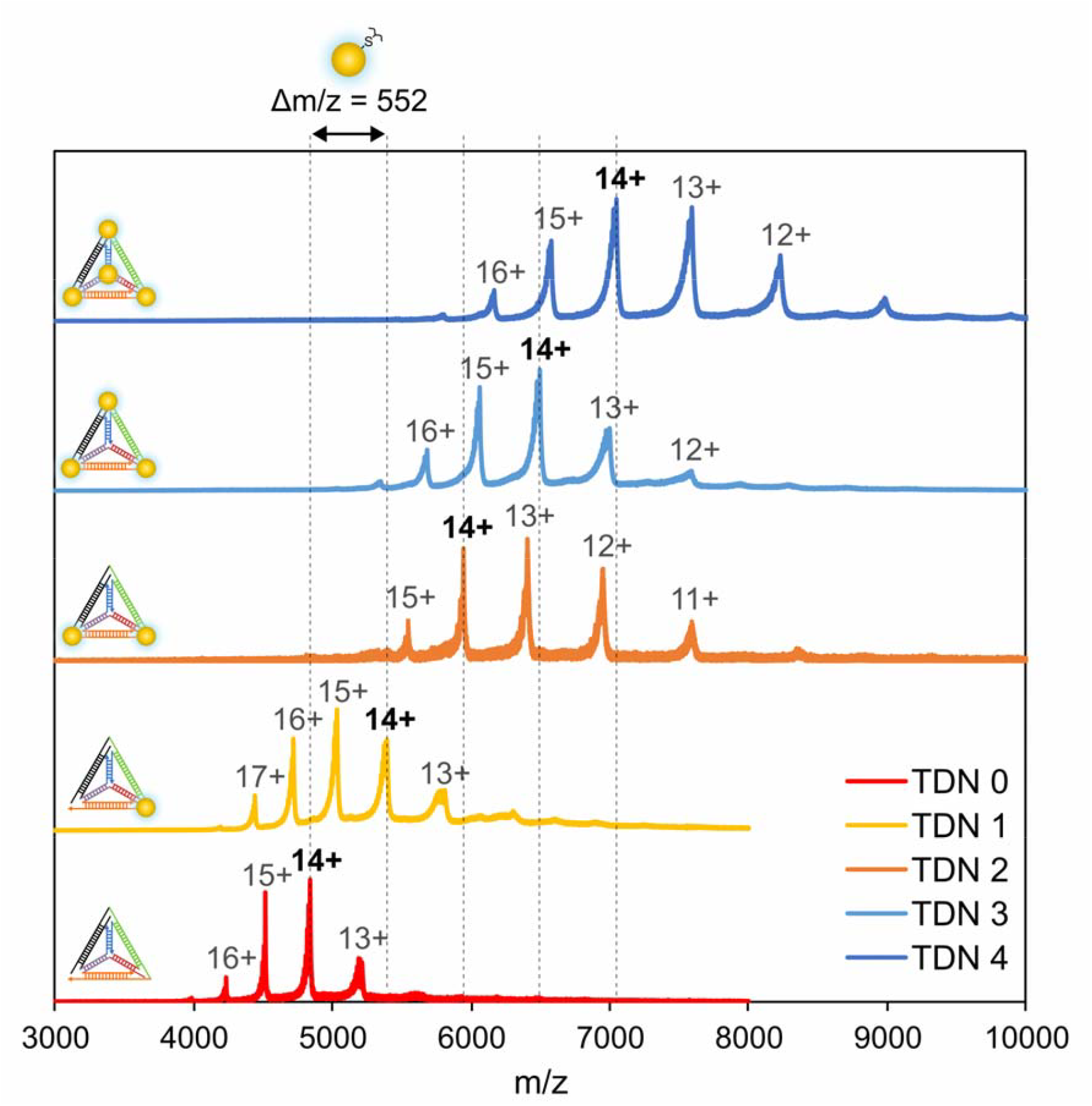
Native MS spectra of the tetrahedral architectures. *m/z* values of the 14+ ions for each TDN are as follow: TDN 0, 4837.25; TDN 1, 5388.76; TDN 2, 5940.69; TDN 3, 6492.24; TDN 4, 7044.59. Each 14+ ion is shifted by a Δ*m/z* of approximately 552, which corresponds to a mass of ―, *i*.*e*., the addition of a gold nanocluster with the thiol function of a ssDNA Au_25_pMBA_17_-S(CH_2_)_6_-PO_3_ (7723 Da).

## IV. Conclusion

A simple method has been developed to synthesize programmable DNA tetrahedra functionalized with a controlled number of atomically precise gold nanoclusters (AuNCs: Au_25_pMBA_18_). Both the ssDNA-AuNCs building blocks and the final DNA tetrahedra were thoroughly analyzed by using a set of complementary techniques. UV-visible and fluorescence spectroscopies were used to validate the presence of both AuNCs and DNA in the samples. Moreover, gel electrophoresis and size exclusion chromatography provided information about the average size and purity of the nanostructures. Additionally, native MS was utilized to gain insight into the accurate composition and stoichiometry of these architectures. The conjugated tetrahedra were determined to be of a high degree of purity, thus validating the efficiency of the complete labelling, assembly and purification processes. Overall, these new biologically guided AuNC assemblies represent powerful platforms for multifunctional applications as theranostics and biophotonic tools.

V.

## Acknowledgements

We acknowledge the support for project MultiMASS from “PTC Instrumentation et Détection” CEA program. X.L.G. would like to thank the French National Research Agency (ANR NanoGold, ANR-22-CE29-0022). D.G. is also grateful to the Labex ARCANE (ANR-11-LABX-003 and CBH-EUR-GS, ANR-17-EURE-0003) that supported part of this study. This work used the MS platform of the Grenoble Instruct-ERIC center (ISBG; UAR 3518 CNRS-CEA-UGA-EMBL) within the Grenoble Partnership for Structural Biology (PSB), supported by FRISBI (ANR-10-INBS-0005-02) and GRAL, financed within the University Grenoble Alpes graduate school CBH-EUR-GS (ANR-17-EURE-0003). The authors would like to thank Laetitia Rapenne for the TEM measurements.

## Notes

### Competing Interest Statement

The authors have declared no competing interest.

