## Supplementary data for "Programmable Assembly of DNA Tetrahedra Bearing Atomically Precise Gold Nanoclusters: Stoichiometric Control and Molecular-Level Characterization"

**Table S1. List of ODN sequences used in this study and structure of the 5'-thiol modifier phosphoramidite monomer**

| Name | Sequence (5' → 3') | Molecular weight (g/mol) |
|---|---|---|
| SS1 | ACA TTC CTA AGT CTG AAA CAT TAC AGC TTG CTA CAC GAG AAG AGC CGC CAT AGT A | 16880.0 |
| SS2 | TAT CAC CAG GCA GTT GAC AGT GTA GCA AGC TGT AAT AGA TGC GAG GGT CCA ATA C | 17032.1 |
| SS3 | TCA ACT GCC TGG TGA TAA AAC GAC ACT ACG TGG GAA TCT ACT ATG GCG GCT CTT C | 16910.0 |
| SS4 | TTC AGA CTT AGG AAT GTG CTT CCC ACG TAG TGT CGT TTG TAT TGG ACC CTC GCA T | 16889.0 |
| 5'SH-SS1 | HS-$(CH_2)_6$-ACA TTC CTA AGT CTG AAA CAT TAC AGC TTG CTA CAC GAG AAG AGC CGC CAT AGT A | 17076.2 |
| 5'SH-SS2 | HS-$(CH_2)_6$-TAT CAC CAG GCA GTT GAC AGT GTA GCA AGC TGT AAT AGA TGC GAG GGT CCA ATA C | 17228.3 |
| 5'SH-SS3 | HS-$(CH_2)_6$-TCA ACT GCC TGG TGA TAA AAC GAC ACT ACG TGG GAA TCT ACT ATG GCG GCT CTT C | 17106.2 |
| 5'SH-SS4 | HS-$(CH_2)_6$-TTC AGA CTT AGG AAT GTG CTT CCC ACG TAG TGT CGT TTG TAT TGG ACC CTC GCA T | 17085.1 |
| 5'-thiol phosphoramidite | 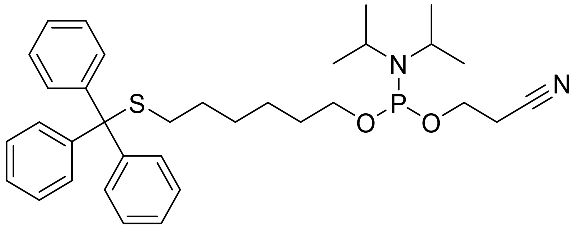 | 576.8 |



**Table S2. Corrected area of the PL emission peaks in relation to the absorbance of the samples at 690 nm with excitation at 690 nm or 808 nm.**

| | $S_{corrected\ 690\ nm}$ (RFU.nm) | $S_{corrected\ 808\ nm}$ (RFU.nm) |
|---|---|---|
| $Au_{25}pMBA_{18}$ | $1.9\times10^5$ | $3.1\times10^5$ |
| $Au_{25}pMBA_{17}SS1_1$ | $3.1\times10^5$ | $4.0\times10^5$ |
| $Au_{25}pMBA_{17}SS2_1$ | $4.8\times10^5$ | $6.4\times10^5$ |
| $Au_{25}pMBA_{17}SS3_1$ | $3.7\times10^5$ | $4.7\times10^5$ |
| $Au_{25}pMBA_{17}SS4_1$ | $3.7\times10^5$ | $4.9\times10^5$ |

**Table S3. Charge states and *m/z* values of the $Au_{25}pMBA_{17}SSX_1$ bricks.**

| | Charge state (*z*) | *m/z* |
|---|---|---|
| $Au_{25}pMBA_{17}SS1_1$ | 8+ | 3076.71 |
| | 7+ | 3515.47 |
| $Au_{25}pMBA_{17}SS2_1$ | 8+ | 3095.76 |
| | 7+ | 3537.07 |
| $Au_{25}pMBA_{17}SS3_1$ | 8+ | 3080.54 |
| | 7+ | 3519.82 |
| $Au_{25}pMBA_{17}SS4_1$ | 8+ | 3077.9 |
| | 7+ | 3516.82 |

These *m/z* values were selected as "starting information" to determine charge states (*z*) and to calculate masses (*m*) of $Au_{25}pMBA_{17}SSX_1$ bricks. The software Masslinks uses these *m/z* values to search for other pairs of *m/z* values. The pairs are used to calculate distinct masses. Then the software calculates an average mass and its error.

**Table S4. Charge states and *m/z* values of the tetrahedral constructs.**

| | Charge state (*z*) | *m/z* |
|---|---|---|
| TDN 0 | 14+ | 4837.67 |
| | 13+ | 5209.90 |
| TDN 1 | 14+ | 5388.61 |
| | 13+ | 5804.35 |
| TDN 2 | 14+ | 5941.18 |
| | 13+ | 6398.20 |
| TDN 3 | 14+ | 6492.26 |
| | 13+ | 6991.59 |
| TDN 4 | 14+ | 7043.20 |
| | 13+ | 7585.34 |

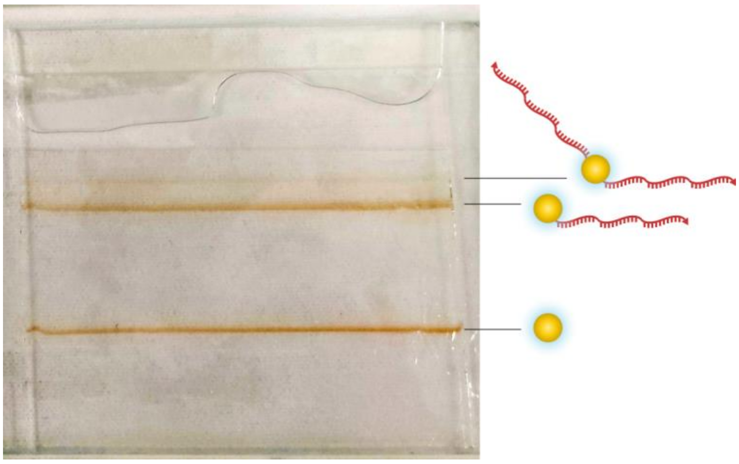

**Figure S1. 20 % 19:1 native polyacrylamide gel electrophoresis (PAGE) analysis of the unpurified $Au_{25}pMBA_{18-n}SSX_n$ in brightfield.**

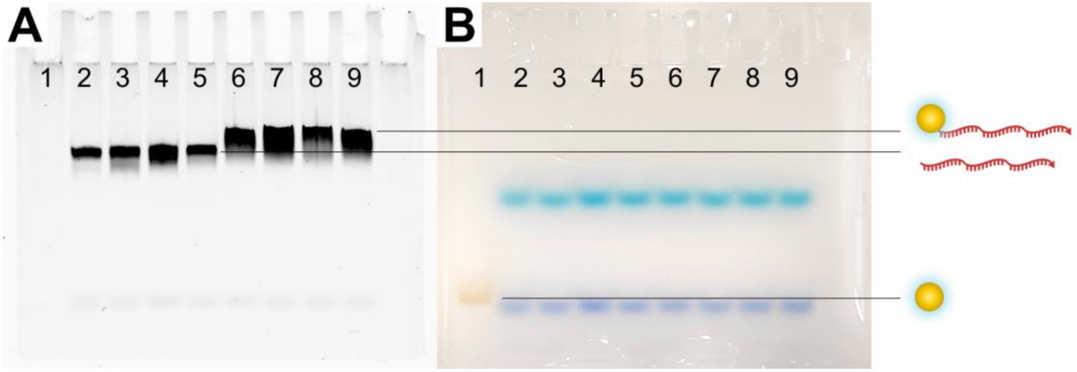


**Figure S2. 20 % 19:1 native polyacrylamide gel electrophoresis (PAGE) analysis of the purified $Au_{25}pMBA_{17}SSX_1$ under UV light (A) and brightfield (B): lane 1, $Au_{25}pMBA_{18}$; lane 2, SS1; lane 3, SS2; lane4, SS3; lane 5, SS4, lane 6, $Au_{25}pMBA_{17}SS1_1$; lane 7, $Au_{25}pMBA_{17}SS2_1$; lane 8, $Au_{25}pMBA_{17}SS3_1$; lane 9, $Au_{25}pMBA_{17}SS4_1$. In the brightfield image, the light blue spots correspond to xylene cyanol and the dark blue spots to bromophenol blue.**

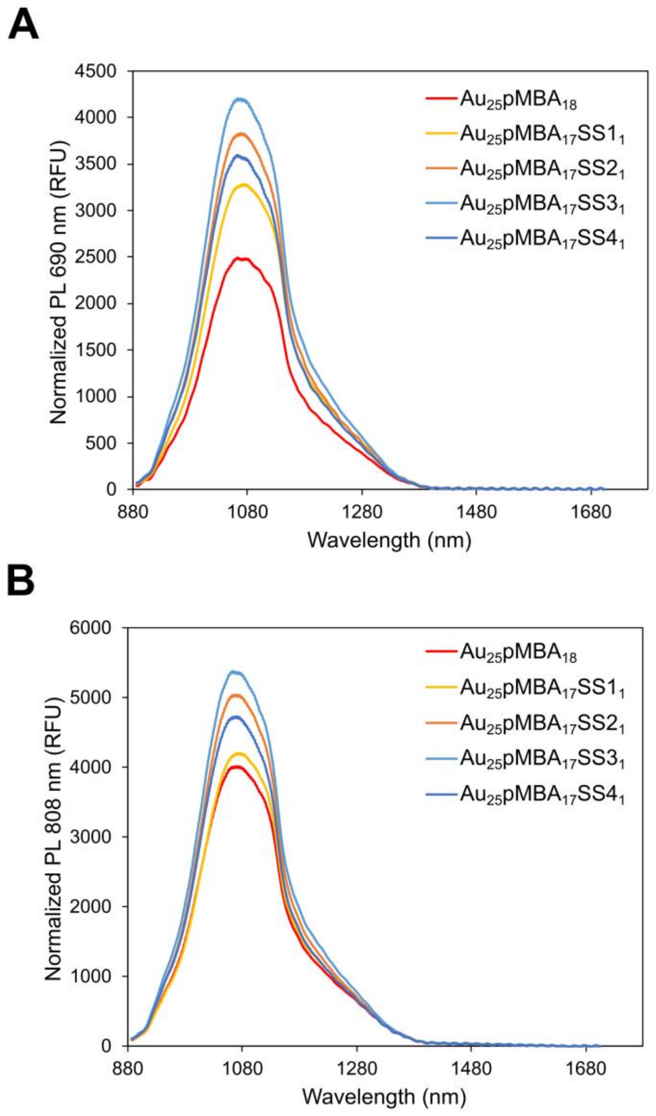


**Figure S3. Photoluminescence (PL) emission peaks of the $Au_{25}pMBA_{18}$ and $Au_{25}pMBA_{17}SSX_1$ constructs with excitation at 690 nm (A) or 808 nm (B).**

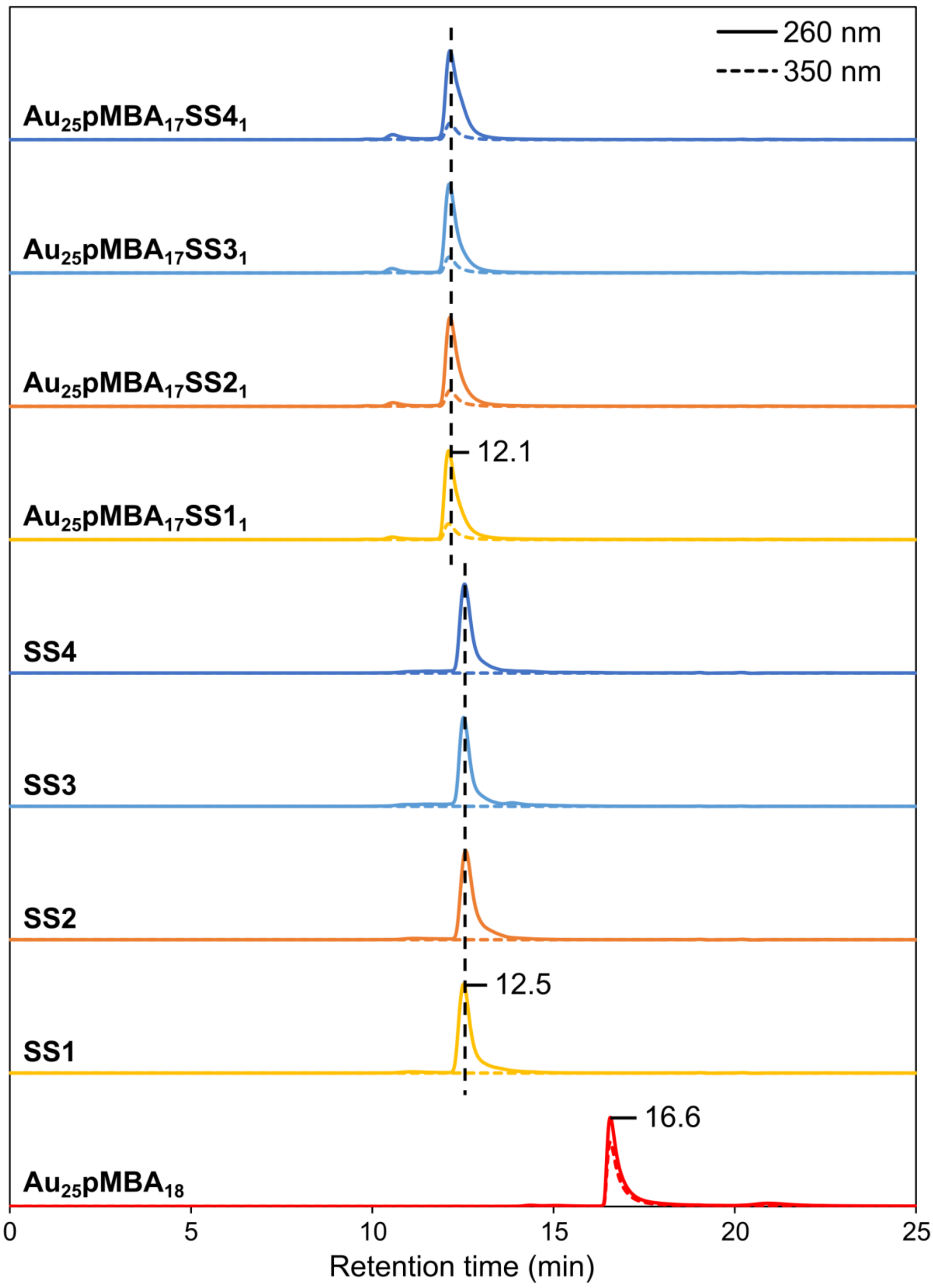


**Figure S4. Size exclusion chromatograms of $Au_{25}pMBA_{17}SSX_1$, SSX and $Au_{25}pMBA_{18}$ with UV detection at 260 nm (solid line) and 350 nm (dotted line).**

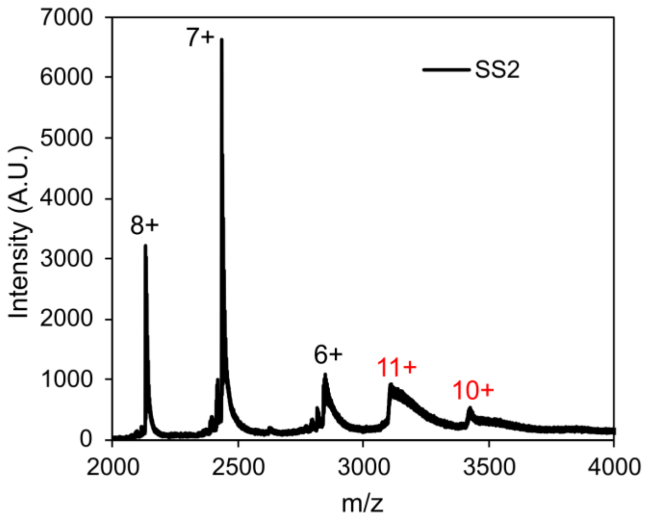


**Figure S5. Native MS spectra of SS2 oligonucleotide. The different charge states of SS2 ions are represented in black while the charge states of SS2 dimer ions are represented in red.**

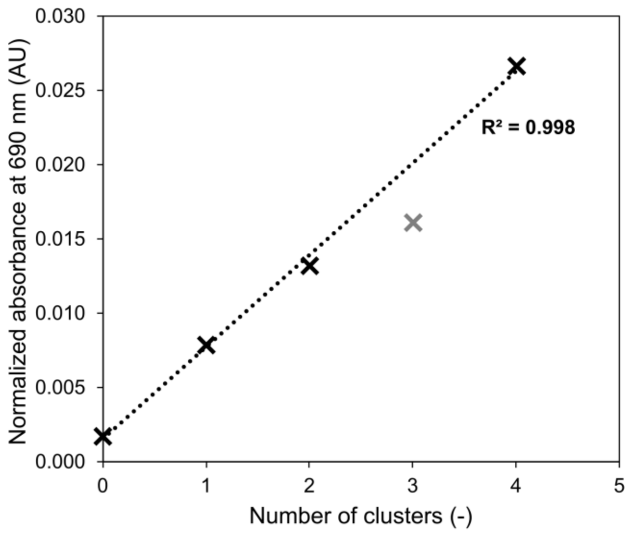


**Figure S6. Normalized absorbance at 690 nm of the different tetrahedra against the number of gold nanoclusters they contain. The data point corresponding to TDN 3 (in gray) was considered an outlier when fitting a linear curve.**

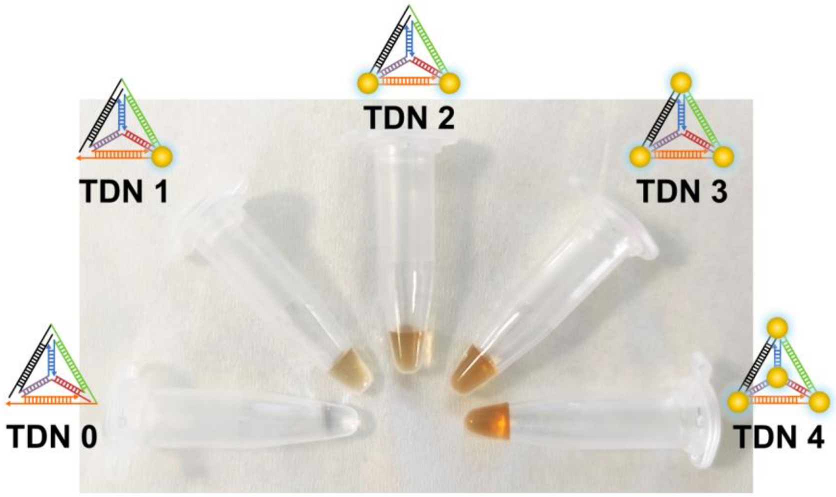


**Figure S7. Picture showing the different tetrahedral solutions at similar DNA concentrations.**

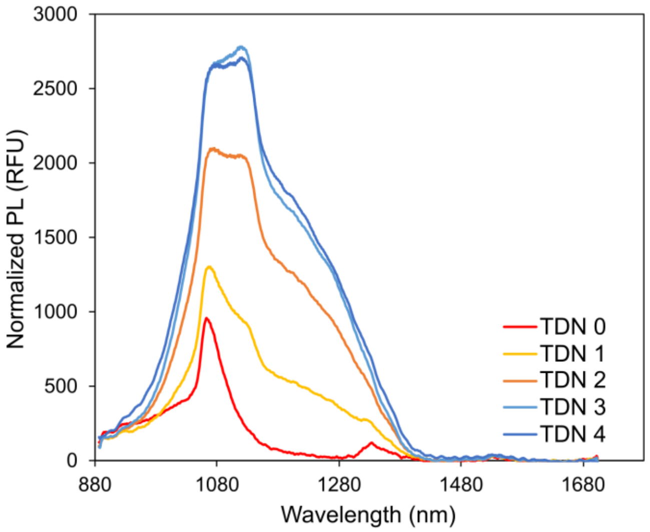


**Figure S8. Photoluminescence (PL) emission peaks of the tetrahedra with excitation at 808 nm.**

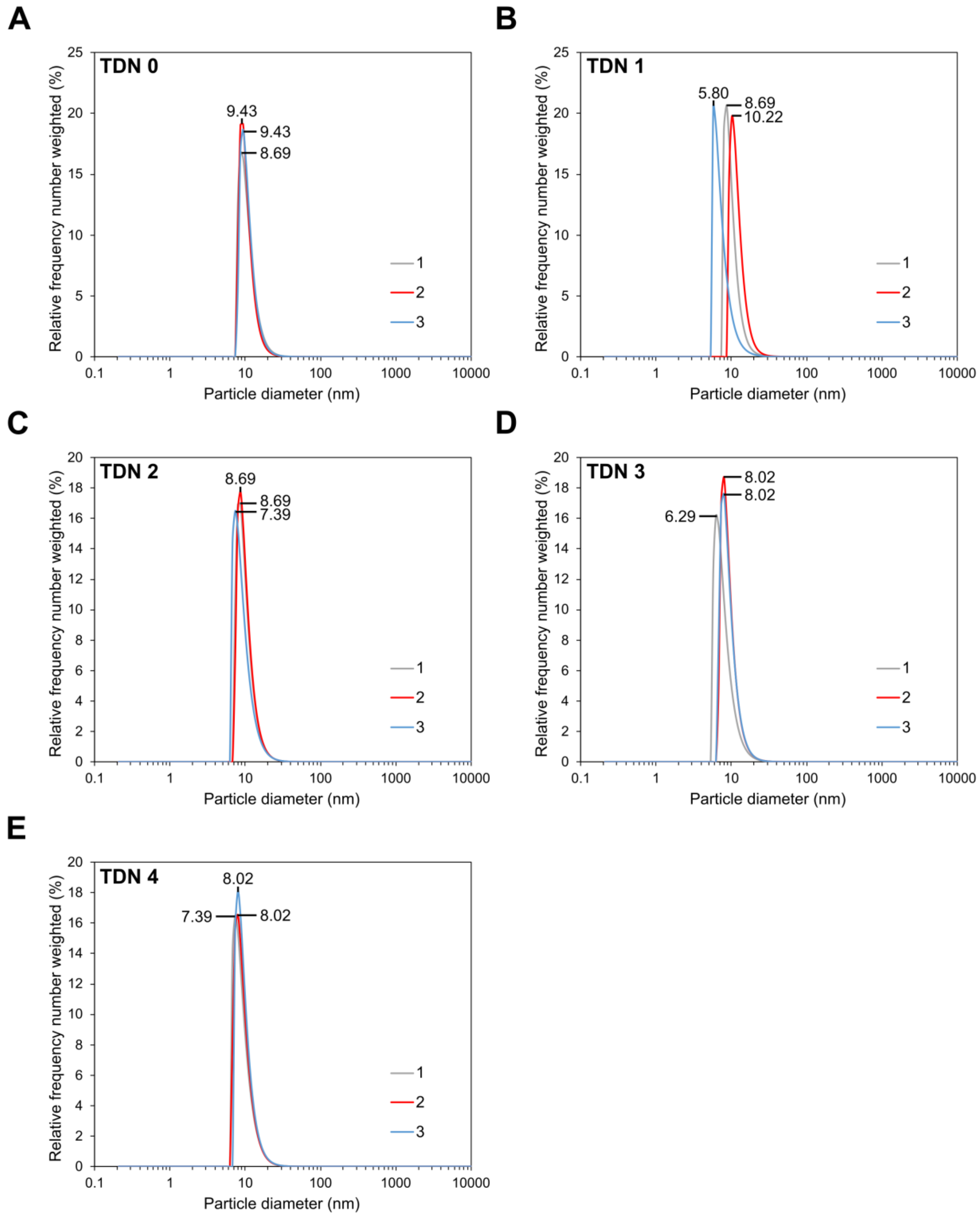


**Figure S9. Number distribution curves of TDN 0 (A), TDN 1 (B), TDN 2 (C), TDN 3 (D) and TDN 4 (E) measured in triplicate using dynamic light scattering (DLS).**

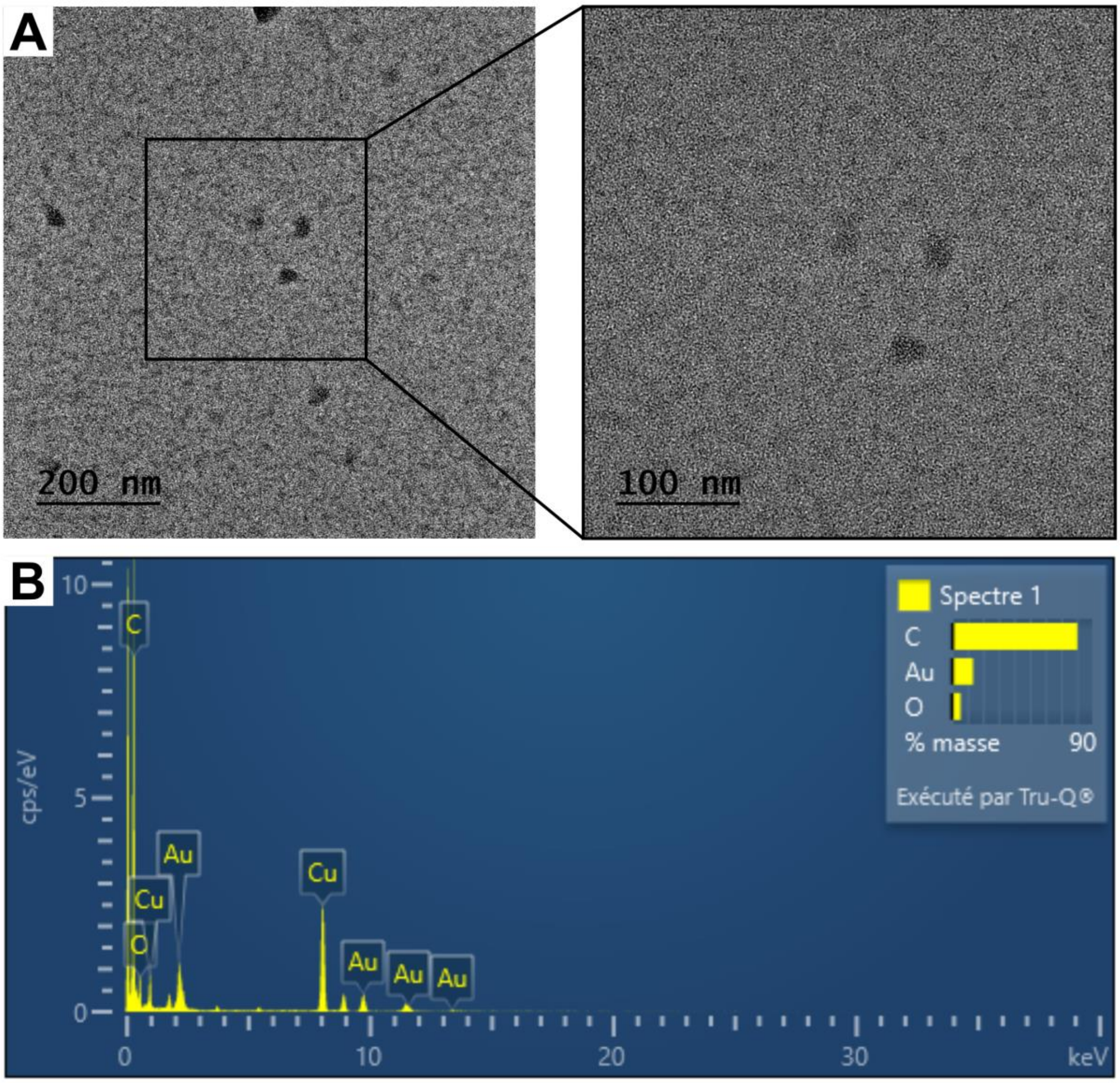


**Figure S10. TEM image (A) and EDX spectrum (B) of TDN 4.**
